# Bacteriocin peer selection for the production of antibiotic selection free biotherapeutic pDNA

**DOI:** 10.1101/2023.10.23.563565

**Authors:** Mohamed El Bakkoury, Luz P. Gómez de Cadiñanos, Philippe Gabant

## Abstract

Plasmid vectors are well established tools used to genetically engineer bacteria both in the laboratory and at industrial scale. The past few decades have seen a rising interest in the use of plasmid DNA (pDNA) for biotherapeutic applications. This interest is a strong driver for the development of technologies to increase pDNA production at biopharmaceutical scale in terms of decreasing production costs and meeting regulatory requirements. Although cell free technologies are emerging, pDNA vectors are still produced by fermentation in *Escherichia coli* strains. As plasmids are extra-chromosomic molecules there is a probability of losing a certain ratio within the *E. coli* population during the fermentation process leading to a decrease of DNA production efficiency. Maintaining pDNA in the population is thus a key element to reach efficient and robust production. Traditionally, antibiotic resistance genes and antibiotics have been used to generate a selective pressure to ensure pDNA stability in the microbial population during the production process. Nowadays, having an antibiotic resistance gene in the pDNA coding sequence represents a limitation both for safety and legal requirements and in terms of production yield. For this reason, we have developed a pDNA antibiotic-free bacteriocin-based selection system, based on the genes involved in the production, processing, secretion and immunity of the bacteriocin microcin V. Our approach is based on the peer pressure exerted by the bacteriocin and does not rely on the addition of any selective agent in the medium to limit population drift and ensure plasmid stability. This novel antibiotic-free approach may be applied to any pDNA vector in different *E. coli* strains and expands their potential applications in both animal and human health as delivery vectors for biotherapeutics.

## 1. Introduction

Since 1973, when Stanley Cohen and collaborators published their work with biologically functioning plasmids *in vitro* (Cohen *et al*., 1973), plasmids have become as one of the most valuable tools in molecular biology, being the workhorse for industrial biotechnology. It is not surprising that for 50 years, cloning genetic functions into a plasmid-based vector has become one of the steppingstones throughout the biological route from research to production. More recently, since the discovery that plasmid DNA injection could be an efficient way to express a specific protein *in vivo* (Wolff *et al*., 1990), the global demand for plasmid DNA (pDNA) has experienced a dramatic increase by the development of the so-called “non-viral” cell and gene therapies, and even more recently by the outbreak of the Covid-19 pandemic. These therapies come in multiple forms but all of them rely at some point, in either their manufacture or mode of action, on the scalable production of pDNA (Ohlson, 2020). Hence, the demand for pDNA will rise leading the scientific community to face important challenges in terms of yield (including cost control) and quality (including regulatory and safety).

However, one of the major drawbacks with using plasmid vectors is their potential segregational instability. The extra-chromosomic nature of these vectors leaves the possibility that during cell division, one of the daughter cells may not receive a plasmid copy, leading to a plasmid-free daughter cell. Under no selective pressure, the plasmid-free population could outgrow the plasmid-bearing population losing the plasmid (Summers, 1991). This genetic drift gives rise to a population that has lost the plasmid and thus the pDNA of interest. To avoid this possibility, the most popular strategy used has been inserting an antibiotic resistance gene in the plasmid and performing antibiotic selection of the host population. In the wild plasmid acquisition mediated by horizontal genetic transfer (HGT) plays an important role not only for adaptation and niche expansion in prokaryotes (Tuller, 2011; Juhas, 2015), but as well as vehicle for resistance gene capture and dissemination (McInnes *et al*., 2020; Che *et al*., 2021). This potential dissemination of resistance genes mediated by industrial pDNA should be limited. It was as early as 1965, when Datta and Kontomichalou showed the widespread transfer of penicillin resistance across Enterobacteriaceae (Datta and Kontomichalou, 1965). Since then, evidence has only increased to the point that, although antibiotics are recognized as one of the greatest achievements of the twentieth century, antibiotic resistance spreading is now considered a global public health emergency. It is now generally accepted that the antimicrobial resistance problem is a complex interconnection between microbes and our planet, the One Health concept. This concept expands the integrative thinking about human and animal medicine, including ecology, public health, and societal aspects (Zinsstag *et al*., 2011). In the case of antimicrobial resistance, the One Health perspective focuses on the risk assessment of emergence, transmission, and maintenance of antimicrobial resistance at the interface between humans, animals, and any linked environment (Robinson *et al*., 2016). Consequently, the application of One Health concept together with the regulatory requirements to which biological agents are subjected are leading towards a “zero-tolerance” rule for antibiotic-based selection in biological production. Hence it is imperative to find alternatives to the use of antibiotic resistance genes and their technological use in vector design, eliminating both the need for antibiotics in bacterial growth media and concerns about the spreading of resistance genes in our ecosystem (Bowater, 2016).

Over the past decades, multiple attempts have been made on the design of antibiotic-free selection systems like those ones involving the complementation of essential genes for example: in an auxotrophic strain, meaning only transformants grow on a defined media lacking the nutrient (González *et al*., 1985), in the operator repressor-titration system where plasmid loss induces downregulation of an essential gene and thus, death (Cranenburgh *et al*., 2001) or with RNA-based approaches that use synthetic or expressed antisense sequences of the target gene (Dryselius *et al*., 2003; Mairhofer *et al*., 2008). Unfortunately, all these systems have in common the potential drawback of requiring the engineering of mutant host strains and/or using specifically defined media. Some strategies have been taken to act after cell division and improve plasmid maintenance, such as the post-segregational killing systems (PSK). The toxin-antitoxin systems (TA) are probably one of the most recognizable PSK mechanisms. Usually in the TA systems plasmid stability relies on the expression of two products: a stable toxin, and an unstable antitoxin, that given its weakness it is not retained in plasmid-free cells upon division and thus, only cells that retain the plasmid can counter-act the toxin and survive (Van Melderen, 2010; Fraikin *et al*., 2020). The successful PSK system of a plasmid-free cell relies on the fact that enough toxin is active in the cell before it goes on to divide again, further diluting the toxin, and that it distributes properly. However, just as with the distribution of plasmids during cell division, there is a giving probability that the distribution of toxin is such that the TA system fails to kill the newly plasmid-free cell leading to a dilution of the plasmid-bearing population, therefore being ineffective during prolonged cultures (Cooper and Heinemann, 2000). When used for example for bioremediation and biotherapeutics applications, plasmids need to be integrated into complex microbial communities were killing the resident species might be undesirable and the objective is just to remove a subpopulation within the community. There have been attempts on controlling populations through *quorum* sensing to control self-killing (Scott *et al*., 2017) or as an alternative to antibiotics against bacterial contamination for example in aquaculture (Ravi *et al*., 2021). However, one of the major weaknesses of the *quorum* sensing control is that it requires the engineering of all strains within the community and in these industrial settings where strains have been optimized for very specific functions this might not be worthwhile. Although these strategies are successful, they are still highly strain dependent, limiting their potential applications.

As an alternative for plasmid vector systems without antibiotics we propose the use of bacteriocins. Bacteriocins represent a huge arsenal of antimicrobial peptides where we can find candidates with the ability to kill closely related strains (narrow *spectrum*) to the bacterial producer or with a broader range of *spectrum* (Cotter *et al*., 2005). A common characteristic for all bacteriocins is the production of an immunity protein, together with the bacteriocin, to ensure survival of the producer bacteria. The immunity genes, which can encode binding proteins, degrading enzymes or export proteins, are usually localized within the same locus as the bacteriocin gene (Smits *et al*., 2020).

We have chosen an adeno-associated viruses-based plasmid (pAAV) where the traditional antibiotic resistance gene has been replaced with either the full operon of a selected bacteriocin, including all the genetic elements required for its production and secretion, or just with the immunity gene of the selected bacteriocin. As proof of concept, we have selected the bacteriocin colicin V present in the PARAGEN collection (Gabant and Borrero, 2019) but it could easily be exchanged for another bacteriocin of interest. Other works have already proven the potential of bacteriocins as PSK mechanisms, such as the Lcn972 system, to stabilize plasmids in *Lactococcus lactis* (Campelo *et al*., 2014) or the use of colicins A and E2 in *E. coli* (Inglis *et al*., 2013). In the same way, colicin V has already been reported to improve plasmid maintenance (Fedorec *et al*., 2019) and the control of a microbial co-culture by the expression and secretion of the bacteriocin against a competitor strain (Fedorec *et al*., 2021). The work presented by Fedorec *et al*., 2019 demonstrated that with traditional TA systems, once plasmid-free cells arise in the population the TA system had no way to save the plasmid-bearing population, whereas colicin V was able to police the entire population due to the toxin being secreted into the environment. This means that when plasmid-free cells arise or if the culture becomes contaminated, the secreted bacteriocin can push the population back toward entirely plasmid bearing leading to peer selection. Nevertheless, all these studies still rely on the use of antibiotics markers for selection, at least for plasmid transformation step, whereas our goal is to remove the antibiotic selection from every step of the process.

For all the above mentioned, hereby we present constructions of vectors that have proven to be stable and efficient pDNA producers without antibiotic selection markers.

Colicin V was the first identified bacteriocin and even though it is still known as “colicin” (Yang and Konisky, 1984; Gilson *et al*., 1987) it should be classified among the microcins (Havarstein *et al*., 1994). Therefore, throughout this paper we will refer to it as microcin V (MccV).

## 2. Results

### 2.1 Construction and performance of a bacteriocin-based plasmid

In this work, we converted the expression vector for recombinant human adeno-associated virus (rAAV) virions, pAAV-RC2 (Cell Biolabs, Inc.), into the plasmid pAAV-MccV as a step further towards the process of efficient antibiotic-free production of plasmid DNA (pDNA).

#### Construction of pAAV-MccV

The plasmid pAAV-MccV was obtained by replacing the ampicillin resistance gene (*Amp*^*R*^) of the commercial plasmid pAAV-RC2 with the complete *mccV* operon (Fragment 1) (Figure 1). The replacement of the *Amp*^*R*^ gene in pAAV-RC2 was achieved by homologous recombination between the R1 and R2 regions of the plasmid with the corresponding regions in Fragment 1 (Figure 1).

**Figure 1.**
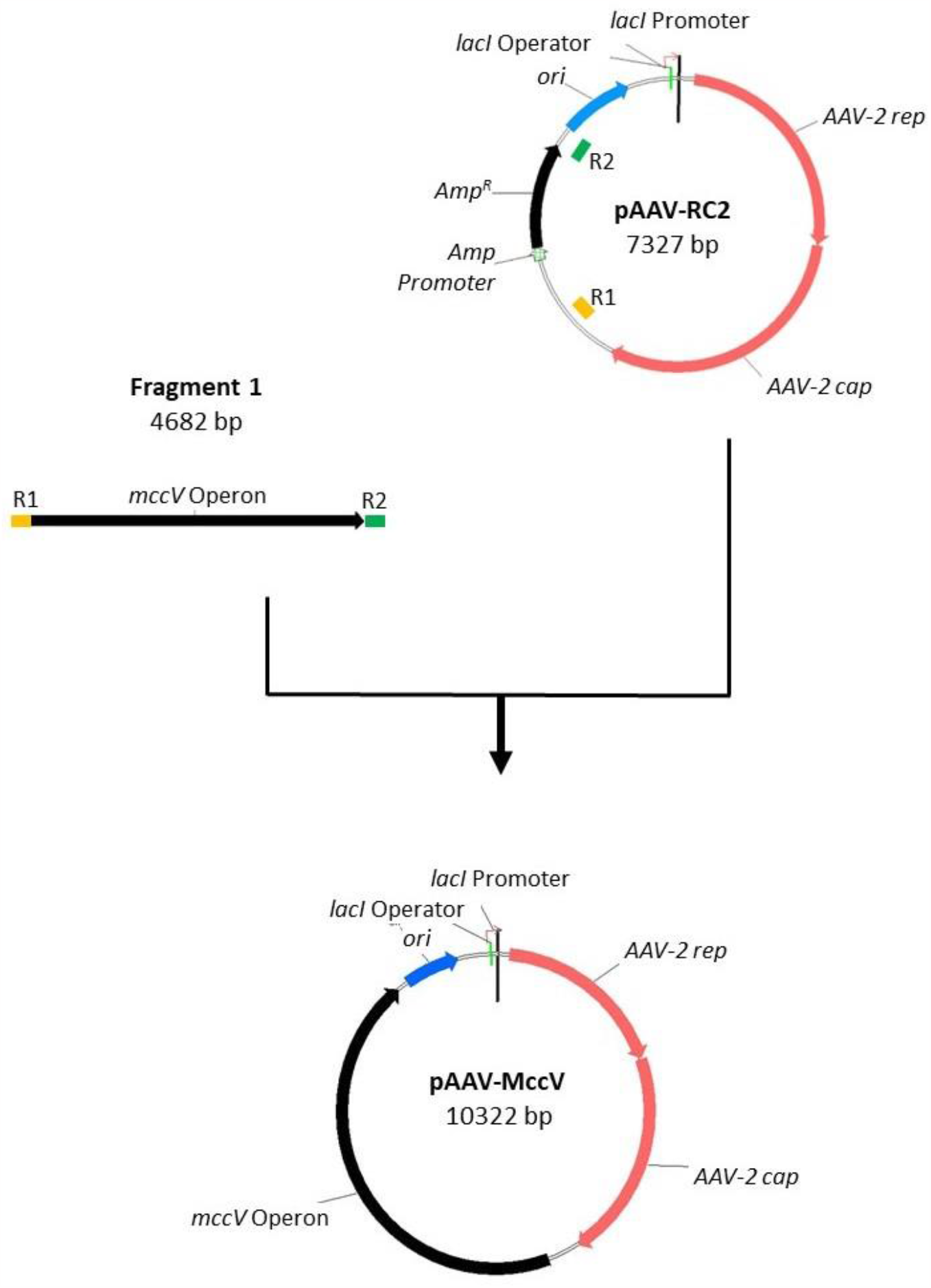
Schematic diagram showing the construction strategy of the plasmid pAAV-MccV. Relevant genes are indicated by arrows whose direction refers to gene transcription. The ampicillin resistant gene (*Amp*^*R*^) from pAAV-RC2 has been replaced by the microcin V operon (*mccV* Operon).

Selection for the plasmid pAAV-MccV was performed by the addition of the bacteriocin MccV to the LB media, although it must be noted that this plasmid not only is immune to the bacteriocin, but it is also able to secrete the MccV to the medium generating a peer selection pressure on the entire population.

#### Growth curve and stability of the pAAV-MccV peer selection vector

In order to measure the effect of bearing the plasmid pAAV-MccV on the host strain *E. coli* STABLE, growth curves were monitored (Figure 2). From the growth curves of the strains STABLE, STABLE (AAV-RC2) and STABLE (AVV-MccV) we observed that the plasmid carrying the microcin MccV operon produces a burden in its host, thus the strain STABLE (AVV-MccV) grew at a significant lower rate (μMax (h^-1^)= 0.85 ±0.23) when compared to the strain STABLE with μMax (h^-1^) of 1.43 ±0.024 and also lower to the strain carrying the commercial plasmid, STABLE (AAV-RC2), with μMax (h^-1^) of 1.15 ±0.14 (Figure 2).

**Figure 2.**
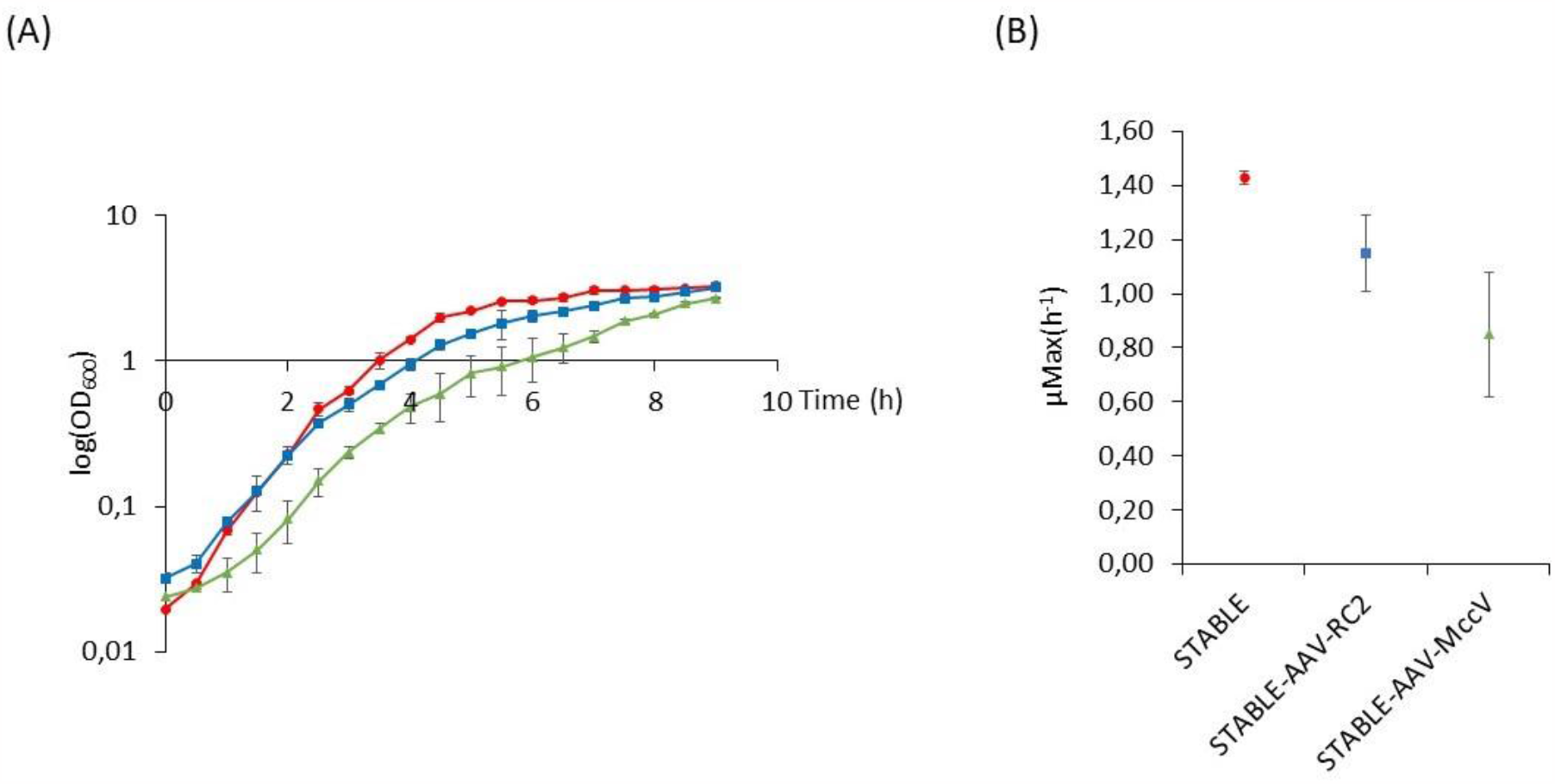
**(A)** Growth curve of the strains *E. coli* STABLE (red), STABLE (AAV-RC2) (blue) and STABLE (AAV-MccV) (green) monitored at 600 nm. **(B)** The average of the estimated maximal growth rates (μMax (h^-1^)) for the three strains are shown.

Plasmid segregational stability within its host cell line was firstly verified after an extended overnight growth (24 h) in non-selective medium followed by a subculture in fresh and non-selective medium until the stationary phase was reached (French and Ward, 1995). For both the control plasmid pAAV-RC2 and pAAV-MccV, we obtained 100% of plasmid retention after 20 hours of growth (Supplementary Material). Secondly, the stability of the plasmid pAAV-MccV within a mixed population was evaluated by following several fresh cultures as shown in Figure 3A: Flasks numbered 1 and 2, with the STABLE (AAV-RC2) strain; flasks 3 and 4, with the STABLE (pAAV-MccV). Flasks 2 and 4 were also inoculated with the plasmid-free STABLE strain, these flasks are thus the challenging culture (Figure 3A). After an overnight incubation we observed an increase of ampicillin-sensitive individuals when analysed the samples from the co-culture between the antibiotic-resistant population and the plasmid-free strain (Flask 2), illustrating the growing advantage of the strain without plasmids. Whereas in the case of the MccV-producing populations (Flask 4) all cells were immune to the bacteriocin and hence able of producing microcin V. Therefore, despite the growing disadvantage of individuals carrying the bacteriocin plasmid, the peer selective pressure generated in the culture broth by MccV was able to rapidly impose individuals harbouring the vector (microcin MccV producer bacteria) and thus eradicating the population of fast-growing plasmid-free bacteria.

**Figure 3.**
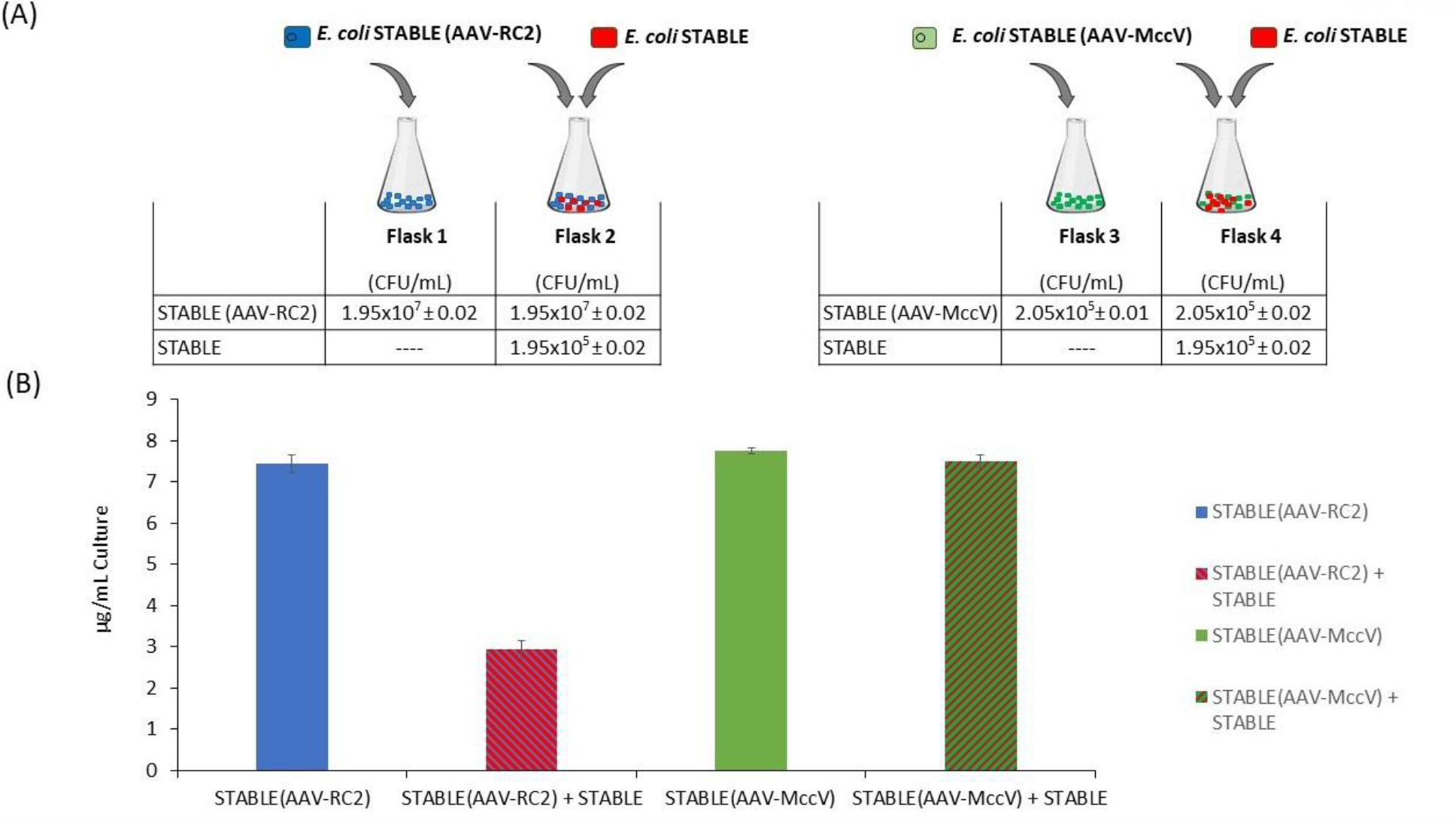
**(A)** Schematic diagram of the different strain combinations stablished: Flask 1 contains the ampicillin resistant *E. coli* STABLE (AAV-RC2) strain (blue); Flask 2, the strain STABLE (AAV-RC2) (blue) and the plasmid-free strain STABLE (red); Flask 3, the STABLE (AAV-MccV) strain (green) and Flask 4, the strain STABLE (AAV-MccV) (green) and the plasmid-free STABLE strain (red). Cell counts at the time of flask inoculation are shown as CFU/mL ± SD for each flask. **(B)** The chart represents the pDNA production obtained from each flask (μg/mL of Culture) after 20 hours of growth.

The above-mentioned results were corroborated when plasmid production was evaluated after an overnight growth (Figure 3B). When the strains STABLE (AAV-RC2) and STABLE (AAV-MccV) were cultured alone (Flasks 1 and 3), similar quantities of pDNA were obtained (7.45 ±0.21 and 7.75 ±0.07 μg/mL of plasmid, respectively). Interestingly, our results also showed that when the strain STABLE (AAV-RC2) was challenged with the plasmid-free strain *E. coli* STABLE (Flask 2), and under no selective pressure, the amount of plasmid recovered was significantly lower than when the strain was cultured alone. Furthermore, the peer selection favoured the STABLE (AAV-MccV) strain when it was equally challenged with the plasmid-free strain (Flask 4) and the amount of pDNA obtained after growing the population without selective pressure was almost the same (7.50 ±0.14 μg/mL of plasmid) as when cultured alone (7.75 ±0.07 μg/mL of plasmid) (Figure 3B). These results are indicative that our antibiotic-free system, thanks to the secretion of MccV, is able to control the peer selection and keep efficiently a population harbouring pDNA.

### 2.2 Plasmid pAAV-ImV, an alternative antibiotic-free system

As our results have shown, strains carrying a pDNA of interest containing a genetic cassette coding for the secretion of the bacteriocin MccV impose a peer selection, allowing the elimination during the production process of bacteria without pDNA copies. This selection is achieved event if we have shown that there is a metabolic cost associated to the production and secretion of MccV. Still, this burden cost is able to efficiently eliminate fast growing plasmid-free bacteria. The use of this peer selection needs to be used for pDNA expected to be difficult to produce since the peer selection is also associated to an increase of vector size which could impact some applications. In order to minimize these effects and facilitate the experimental handling throughout the production process of pDNA, we also constructed and tested a second plasmid, the pAAV-ImV, which does not express the bacteriocin but harbours its immunity gene (Figure 4). This vector is only ≈6 kb and thus much smaller compared to the pAAV-MccV which is around 10 kb, making this construction more compact even than the plasmid used as backbone, the commercial and antibiotic resistant pAAV-RC2 (≈7 kb) (Figures 1 and 4).

**Figure 4.**
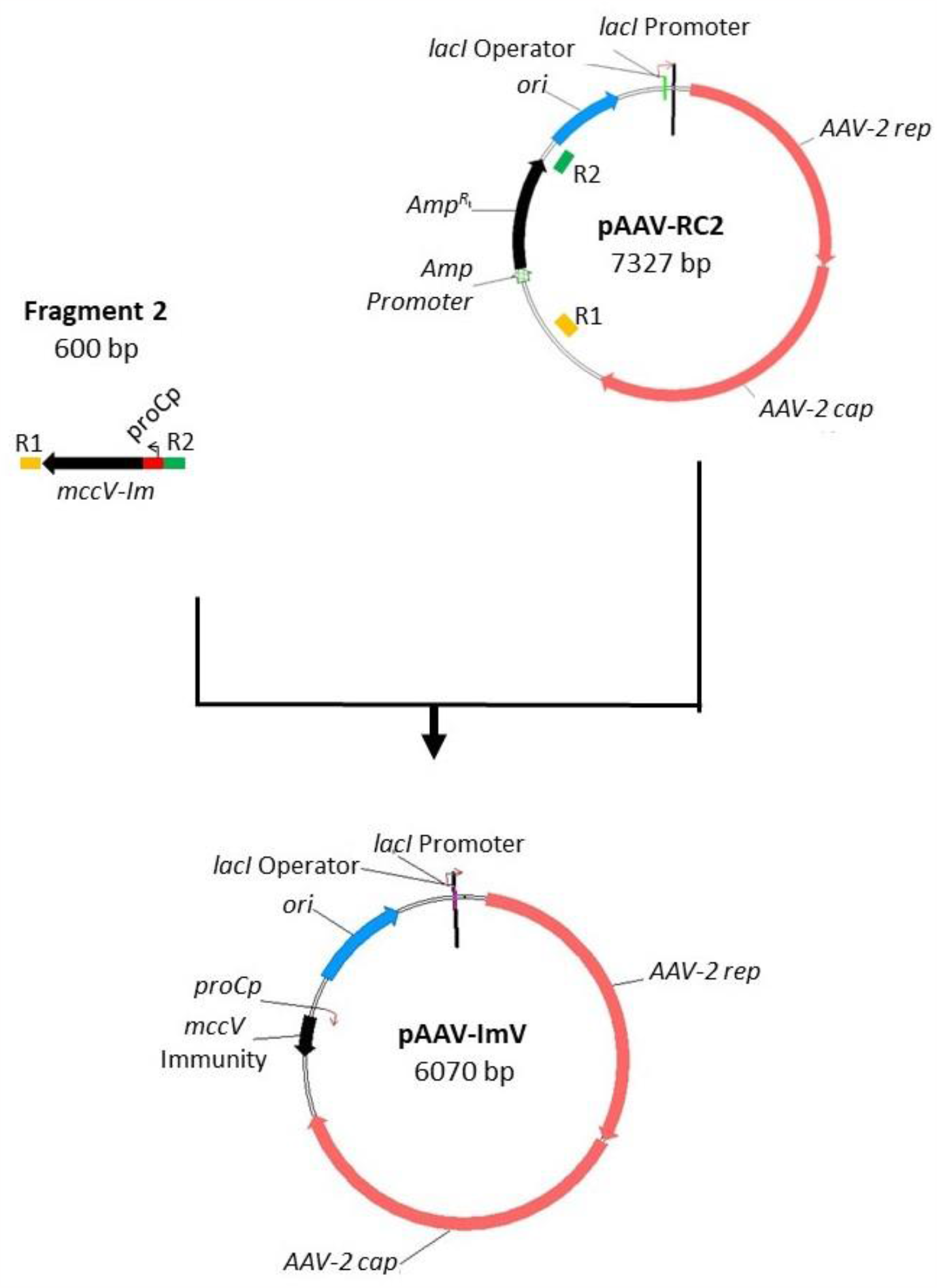
Schematic diagram showing the construction strategy of the plasmid pAAV-ImV. Relevant genes are indicated by arrows whose direction refers to gene transcription. The ampicillin resistant gene (*Amp*^*R*^) from pAAV-RC2 has been replaced by the immunity gene of microcin V (*mccV Immunity*).

The construction of the plasmid pAAV-ImV was achieved by swapping the ampicillin resistance gene (*Amp*^*R*^) of the commercial plasmid pAAV-RC2 with the immunity gene of microcin MccV (*mccV* immunity) (Fragment 2) (Figure 4). The expression of the *mccV immunity* gene in pAAV-ImV is under the control of the *proCp* promoter (Davis et al., 2011) carried in Fragment 2 (Figure 4). The replacement of the *Amp*^*R*^ gene in pAAV-RC2 was achieved by homologous recombination between the R1 and R2 regions of the plasmid with the corresponding regions in Fragment 2 (Figure 4).

### Performance of plasmid pAAV-ImV

We observed that the strain harbouring the pAAV-ImV plasmid presents the same growing fitness than the initial vector, showing that the burden observed with pAAV-MccV is well generated by the production and secretion the bacteriocin MccV used for peer selection. Hence, the strain STABLE (AAV-ImV) was able to achieve a maximal growth rate (μMax (h^-1^)) closer to the wild-type strain (1.204 ±0.017 and 1.430 ±0.024, respectively) and significantly higher than the STABLE-AAV-MccV strain (0.85 ±0.23). After growing for an overnight, the pDNA obtained from the strain STABLE-AAV-ImV was comparable to the quantity obtained from the commercially optimized plasmid pAAV-RC2 with no significative difference between them (Figure 5). The stability of pAAV-ImV was tested after an extended overnight growth in non-selective medium. For both plasmids, the pAAV-ImV and the control plasmid pAAV-RC2, we obtained 100% of plasmid stability (Supplementary Material).

**Figure 5.**
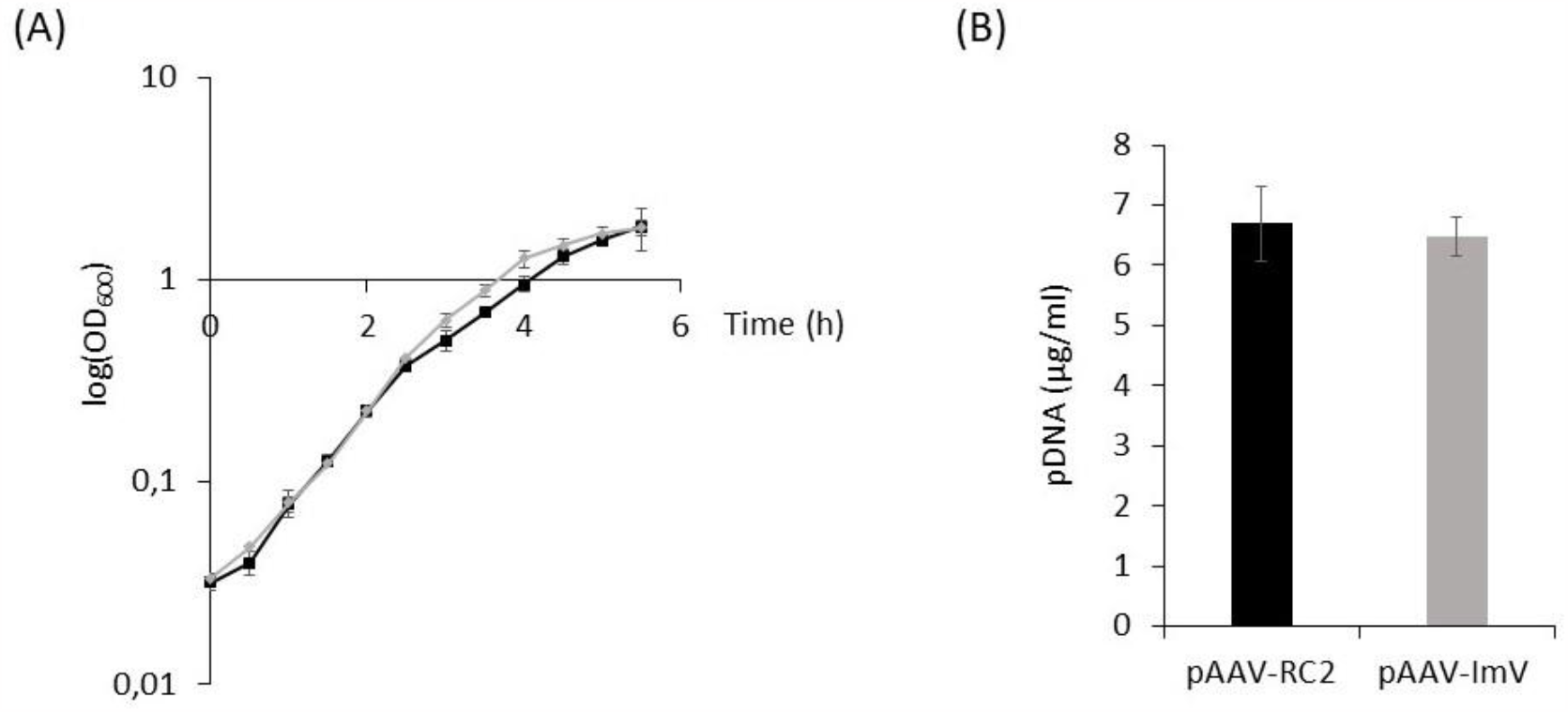
**(A)** Growth curves of the strains *E. coli* STABLE (AAV-RC2) (in black, ▪) and STABLE (AAV-ImV) (in grey, 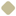) monitored at 600 nm. **(B)** pDNA Quantification after extraction of plasmids pAAV-RC2 and pAAV-ImV.

### 2.3 Measurement of the plasmids superhelical densities

Production of pDNA that may be used for biotherapeutic applications requires high quality of the pDNA. The quality of the pDNA is measured by percentage of supercoiling and the supercoiling density as plasmids in their supercoiled isoform are considered to have a superior stability and biological activity in comparison to other isoforms (Hassan *et al*., 2016). Based on this criterion, we determined the superhelical density of both of our plasmids, pAAV-MccV and pAAV-Im and compared them to the one commercially optimized vector pAAV-RC2. Figure 6 shows a sample of two-dimensional and one-dimensional chloroquine agarose gels which were run to determine the linking difference of the plasmids. Each sample was run on a single gel and each band represents the different level of supercoil form that is found on each plasmid. Interestingly, for the smaller plasmids, the pAAV-ImV and the control pAAV-RC2, the two-dimension gels show the nicked form of the molecules and their transition from a relaxed configuration to being positively supercoiled by the effect of the *E. coli* topoisomerase I. Whereas in the case of plasmid pAAV-MccV, due to its higher size (≈10 kb), we were able to observe the full migration of the topoisomers: the nicked forms and from negatively to positively supercoiled. For an extended understanding on the superhelical plasmid determination through two-dimensional gels, please refer to Gibson *et al*., 2020 (Gibson *et al*., 2020).

**Figure 6.**
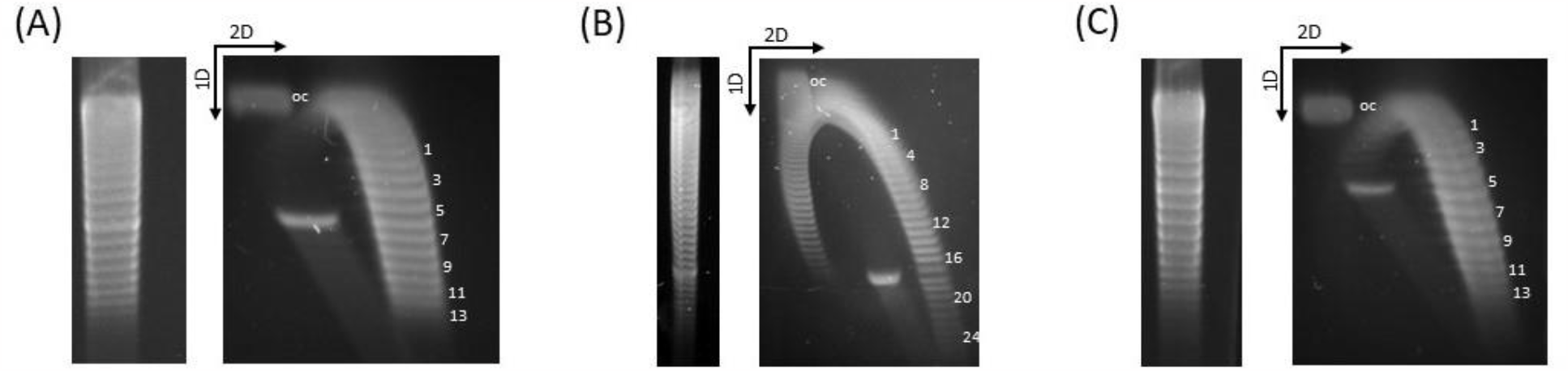
Superhelical density determination of plasmids: **(A)** control plasmid pAAV-RC2, **(B)** pAAV-MccV and **(C)** pAAV-ImV. The 2D and 1D chloroquine-agarose gel electrophoresis show the distribution of topoisomers of the plasmids. The bands depict the linking difference which is then used to calculate the superhelical density (σ). OC, Open-circle form.

The linking difference with the linking number was used to calculate the superhelical density (σ) of the plasmid according to the equations (1 to 3) described in the materials and methods section of this manuscript. The result of the comparison of the superhelical density between the plasmids (pAAV-RC2, pAAV-MccV and pAAV-ImV) amplified in the three *E. coli* strains STABLE (AAV-RC2), STABLE (AAV-MccV) and STABLE (AAV-ImV), respectively, was expressed as the mean ±SD (Figure 7A). As shown in the chart, no statistical difference was observed between the average superhelical density of the plasmids. As it may be observed in Figure 7B, all three plasmid preparations successfully achieved 95% of the supercoiled form, being only the remaining 5% OC forms.

**Figure 7.**
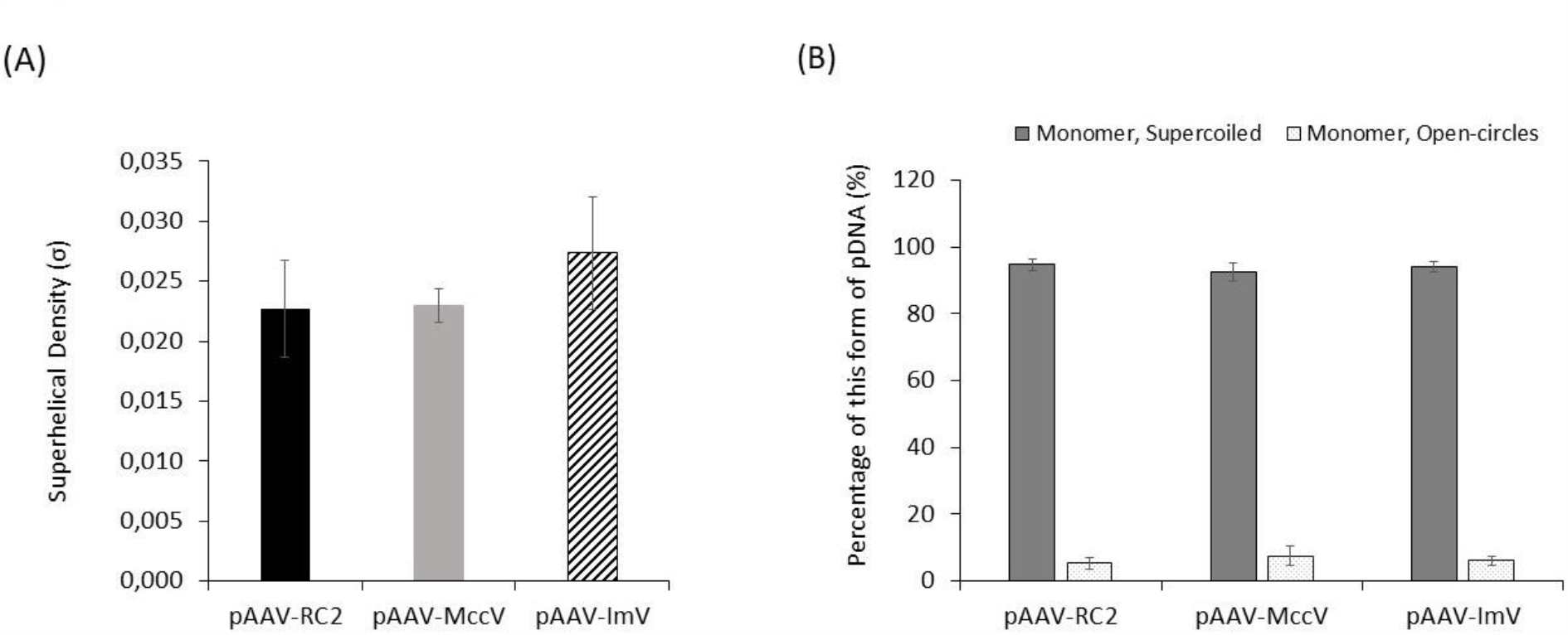
**(A)** Average superhelical density of the plasmids pAAV-RC2 (black), pAAV-MccV (grey) and pAAV-ImV (stripes), n=9. **(B)** The average of total percent supercoiling and open-circle topoisomers of the different pDNA preparations are shown.

These results demonstrate that efficient and high quality pDNA production is possible with stable and antibiotic-free plasmids like pAAV-MccV and pAAV-ImV.

## 3. Discussion

There is an urgent need for alternative systems to antibiotic resistance genes for vectorial plasmid selection in industrial production. This is required to make production more cost-effective, robust, and flexible as well as in line with regulatory rules that are imposing a zero-tolerance for the use of antibiotics in bioprocesses. The exploration of new selection markers is also driven by the need of new developments in the field of synthetic biology applied to precision fermentation. These innovations required in selection developments are imposed by the fact that microbes are prompt to evolve and so frequently drift from their initial design and by doing so they generate considerable industrial challenges in terms of scaling up. Because of their own nature, bacteriocins *loci* represent a unique technological alternative to antibiotics and their resistance genes for any in precision fermentation including the plasmid DNA (pDNA) production vectors used in biotherapeutic applications. This potential has been already identified for plasmid stabilization in *E. coli* but so far in combination with antibiotic markers.

The current strong need for pDNA and the foreseen need to produce these vectors for innovative DNA-based vaccination, gene therapy and also animal health, has motivated us to design new vectors without the antibiotic resistance genes. Based on published results of Fedorec *et al*. 2019 & 2021 (Fedorec et al., 2019, 2021) on the potential of the bacteriocin microcin V (MccV) to ensure plasmid stability in *E. coli*, we decided to base our work on this bacteriocin locus with the aim of building antibiotic-free pDNA vectors. Due to the importance of vectors derived from adeno-associated viruses (AAVs) in therapy (Zinn and Vandenberghe, 2014; Shupe *et al*., 2022), we decided to use as plasmid backbone the commercially available pAAV-RC2 (Cell Biolabs, Inc.). Our approach consisted in replacing this vector’s ampicillin cassette with in one hand, the full operon of MccV which led to the plasmid pAAV-MccV, and on the other hand with just the immunity gene of MccV (*mccV-Im*) which led to the vector pAAV-ImV.

The two antibiotic-free generated constructs were tested for their impact on host growing behaviour. We observed that the production of MccV for peer selection has a metabolic cost, impacting the growth rate of *E. coli*. By stablishing competition experiments we were able to show that even slower growing strains containing the peer selection circuit were able to outcompete the faster growing bacteria, the ones without any vector, leading to a 100% bacteriocin-resistant culture. This result was somehow unexpected, as the experiments were performed without using any selective pressure in the media, therefore it would be expected that the plasmid free strain would outcompete the bacteriocin producer strain (De Gelder *et al*., 2007) as it did happen with the antibiotic-resistant strain. Our results demonstrate that this “predation” phenotypes of the cells harbouring the pDNA of interest was very efficient and stable. We observed no differences in terms of pDNA production between culturing the bacteriocin-producing strain alone or in a co-culture with a plasmid-free strain, whereas in the case of the antibiotic resistant strain there was a 60% loss in pDNA production when it was co-cultured compared to when it was alone. Meaning that the production and secretion of bacteriocins to the environment confers the plasmid-bearing population a clear peer selection advantage over other close related contaminant bacteria that might appear during fermentations for pDNA production.

Nevertheless, having the full operon of a bacteriocin, in this case MccV, in the pDNA producing vector might not be necessary for all processes or applications and it does come out at the cost of increasing the vector size, with the impact that might have in the process. A shorter plasmid size, around 6 kb, is always desirable so that the target insert may include larger sequences, such as genomic *loci* or more than one therapeutic gene in the same vector (Lara and Ramírez, 2012). Therefore, for those cases where peer selection is not essential, we designed and constructed an alternative system: the vector pAAV-ImV. This vector has a reduced size of ≈6 kb and carries the immunity gene of MccV as selection marker, without the need of antibiotics. Our experimental results showed that losing the capability of producing and secreting the bacteriocin to the extracellular medium, lifted the energy burden and the host strain carrying the pAAV-ImV had a growth rate comparable to the wild-type strain.

The plasmid constructions presented in this work were also analysed and compared to the commercial plasmid in terms of quality. Supercoiling is essential for the application of pDNA with clinical purposes, as it is believed that this form is better protected against enzymatic degradation and thus more efficient as more plasmid can reach the nucleus of the target cell *(Cupillard et al*., *2005; Pillai et al*., 2008; Baranello *et al*., 2012). DNA supercoiling refers to the degree of unwinding of a DNA duplex. A “relaxed” double-helical segment of DNA consists of two DNA strands twisting around the helical axis once every 10.4–10.5 sequence base pairs (Hassan *et al*., 2016). Our results demonstrate that both our vector pAAV-MccV and pAAV-ImV preparations, achieved ≈95 % of supercoiled content just like the optimized commercial plasmid and in compliance with the USA-FDA recommendations (Hitchcock *et al*., 2010).

Throughout this manuscript we have demonstrated that it is possible to achieve an efficient and high quality pDNA production with antibiotic-free plasmids, but we believe that there is also a need for strain design. The experimental work presented here has been performed using the standard strain STABLE due to its convenience for our experimental work. However, we sequenced the strain’s genome with our MinION-101B sequencer (Oxford Nanopore Technologies plc.) and learnt that the commercially available *E. coli* STABLE strain not only has an aggregation phenotype and harbours the F1 plasmid but also carries a chromosomal mutation that confers resistance to the antibiotic streptomycin. Hence, although it has served its purpose as host strain for demonstrating the advantages of the vectors pAAV-MccV and pAAV-ImV over conventional plasmids, we believe that strain design is mandatory in order to go further towards antibiotic-free pDNA production. For this reason, in parallel to the work presented here, we have worked on “re-shaping” the STABLE strain by removing the F1 plasmid and the chromosomal mutation that leads to antibiotic resistance (manuscript in preparation).

The work presented here, using the bacteriocin MccV as antimicrobial agent for pDNA production, serves as proof of concept for this technology. Any other bacteriocin of choice could be specifically selected depending on the application or the contaminant(s) of interest, thus opening the door to an almost limitless number of tailor-made vectors. We are strongly convinced that bacteriocins have the potential to be alternative sources of antimicrobial agents in the fight against AMR. It should be noted that the bacteriocin nisin, for example, has been used for decades, yet widespread resistance has not been reported. Furthermore, there is an increasing trend on testing bacteriocins *in vivo* models to prove their lack of toxicity (Benítez-Chao *et al*., 2021).

Like most other fields, the use of bacteriocins also has problems to address, like the cost of their large production processes but the incorporation of synthetic biology into their pipeline of production is a way forward and they are certainly, as this work demonstrates, a valuable tool for a further step towards total antibiotic-free pDNA production processes.

## 4. Conclusions

In this publication we exemplify, using microcin V for the first time, that extra-chromosomic plasmid vectors can be constructed and amplified without antibiotic resistance genes. Our competition experiments have demonstrated that a peer-based selection represents a burden in terms of growth rate, but the selection imposed by the secretion of the bacteriocin is so strong that allows to overcome the genetic drift. From all the applications for which these technologies could be of use, we decided to focus this research on pDNA production. The results presented here confirm that antibiotic-free plasmid vectors based on bacteriocins are a high quality efficient pDNA production systems. Further experiments are required, but we believe that the present study lays the groundwork for increasing scalability of future cell factories through stable genetic designs that sustain a high-production phenotype and enable long-term production.

## 5. Materials and Methods

### 5.1 Bacterial strains, constructions, and plasmids

All microorganisms, plasmids and genetic constructs used during this study are listed in Table 1. The original wild type producing strain for bacteriocin microcin V (MccV) was a gift from Prof. Chris P. Barnes and his team from University College London (UK). For the experimental work described in this paper we synthetized the complete *mccV* operon. Competent *Escherichia coli* STABLE cells (New England Biolabs) were used as host strains for recovering the recombinant vector. Strains were grown at 37 °C with vigorous agitation in Lysogeny Broth (LB) (Sambrook and Russell, 2001) and transformants were recovered in LB agar plates containing a suitable concentration of bacteriocin or antibiotic when needed. All transformations were achieved by using standard protocols for *E. coli* strains (Hanahan, 1983).

**Table 1.** *Escherichia coli* strains and plasmids used in this study.

| Strains, Constructions & Plasmids | Relevant Characteristic(s) | Source |
| --- | --- | --- |
| **Strains** |  |  |
| STABLE | *E. coli* strain used for plasmid DNA production. | NEB |
| STABLE (AAV-RC2) | *E. coli* STABLE transformant carrying plasmid pAAV-RC2. | This study |
| STABLE (AAV-MccV) | *E. coli* STABLE transformant carrying plasmid pAAV-MccV. | This study |
| STABLE (AAV-ImV) | *E. coli* STABLE transformant carrying plasmid pAAV-ImV. | This study |
| **Constructions** |  |  |
| Fragment 1 | *In vitro* synthesized DNA fragment (4682 bp) comprising the microcin MccV operon (*mccV*) flanked by the R1 and R2 regions homologous to the pAAV-RC2 vector. | This study |
| Fragment 2 | *In vitro* synthesized DNA fragment (600 bp) comprising the immunity gene of microcin MccV (*MccV-Im*) under the control of proCp promoter flanked by the R1 and R2 regions homologous to the pAAV-RC2 vector. | This study |
| **Plasmids** |  |  |
| pAAV-RC2 | Expression vector for recombinant human adeno-associated virus (rAAV) virions; *Amp<sup>R</sup>*. | Cell Biolabs, Inc. |
| pAAV-MccV | pAAV-RC2 derivative with *Amp<sup>R</sup>* replaced by *mccV* operon. | This study |
| pAAV-ImV | pAAV-RC2 derivative with *Amp<sup>R</sup>* replaced by *mccV* immunity gene. | This study |

NEB, New England Biolabs; *Amp<sup>R</sup>*, ampicillin resistance gene.

The inserts used for the construction of the desired plasmids were synthesized by GENEWIZ (Azenta Life Sciences, Germany), prepared by polymerase chain reaction (PCR) amplification followed by gel purification. PCR primers are provided in Table 2. Vectors were digested with restriction enzymes (New England Biolabs) and following digestion, both insert and vector were gel-purified using MicroElute Gel Extraction Kit (Omega Bio-Tek, Inc.) following manufacturer’s instructions prior to ligation. Some recombinants were generated by homologous recombination using GenBuilder Kit (GenScript Biotech Corporation) following manufacturer’s instructions.

**Table 2.**
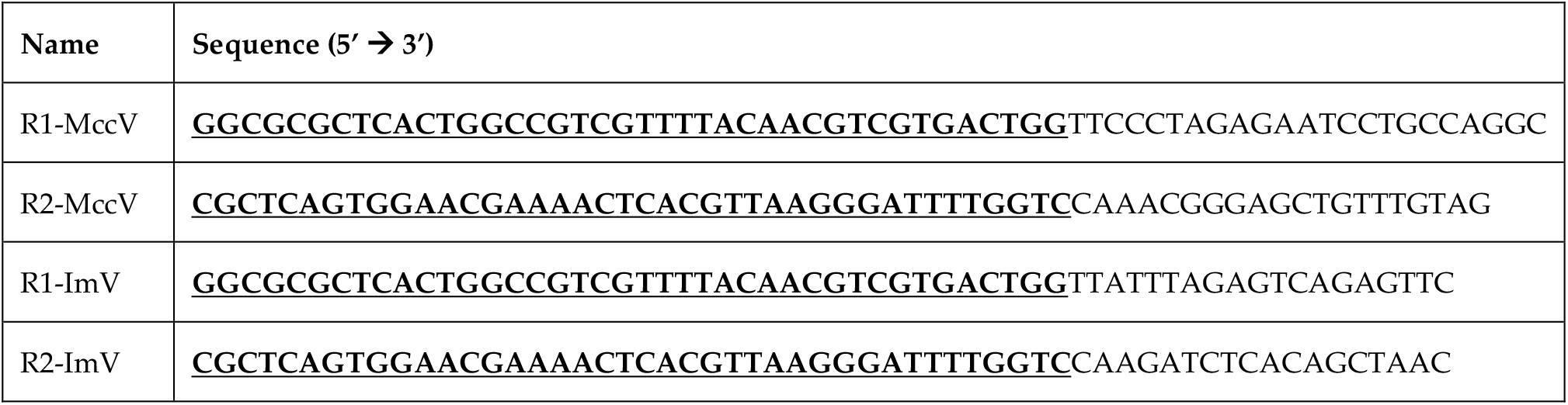
Primers used for vector constructions. Underlined bold nucleotides indicate the sequence for homologous recombination.

### 5.2 Growth curves and Maximal growth rate

Optical density (OD) measurements at 600 nm were used to follow the growth of the different strains in LB broth during time and to estimate their maximal growth rate (μMax). The estimated value for μMax(h^-1^) was calculated by fitting the curves to a sigmoid model using the Microsoft Excel add-in DMfit v.3.5 (available at https://www.combase.cc/index.php/en/).

### 5.3 Drift control assay

Plasmid DNA (pDNA) stability was tested by setting a drift control assay carried out with different co-cultures of *E. coli* strains: the wild-type, STABLE; the ampicillin resistant, STABLE (AAV-RC2) and the microcin V producer, STABLE (AAV-MccV) (Table 1).

A 50-fold-diluted fresh culture from each overnight seed culture was grown to exponential phase. Once the desired cell population of the three strains (STABLE, STABLE (AAV-RC2), STABLE (AAV-MccV)) was reached, four different flasks with LB medium were inoculated. At the time of inoculation, a sample from each culture was taken and spread onto agar plates (1.5%) for CFU (colony forming unit) counting. After an overnight incubation with vigorous shaking at 37 °C, samples were taken for CFU counting, phenotyping as well as pDNA production was assessed by pDNA extraction and quantified. Viability counts were performed after spreading the samples onto non-selective (LB) and selective agar plates for CFU counting and phenotype screening. Selective agar plates consisted, on LB-Amp (100 μg/mL ampicillin) for samples involving the strain STABLE-AAV-RC2, and LB with the microcin MccV (LB-MccV) (50-fold-diluted MccV, Minimum Inhibitory Concentration (MIC) of 1/100) for samples with the strain STABLE (AAV-MccV).

### 5.4 Plasmid DNA maintenance and quantification

The maintenance of each plasmid within its host cell line was verified by inoculating a competent *E. coli* STABLE strain with each of the plasmids used throughout this work (pAAV-RC2, pAAV-MccV and pAAV-ImV). After an extended overnight incubation (24 h) in non-selective LB medium at 37 C with vigorous shaking, a diluted sample from each culture was used as fresh inoculum of LB medium and let grow until the stationary phase was reached. At this point, a sample from each cell line was spread onto selective and non-selective agar plates. The percentage of plasmid retaining cells was determined from the ratio of colonies on selective plates over those on non-selective plates. The experiment was done three times with triplicate samples from each cell line (n=9).

The pDNA production obtained with the different strains was evaluated by extracting the pDNA with the MicroElute Gel Extraction Kit (Omega Bio-Tek, Inc.) and quantifying it with Qubit Invitrogen (ThermoFisher Scientific). As quality control, the plasmid extractions were analyzed by gel electrophoresis after enzymatic restriction.

### 5.5 Determination of plasmid superhelical density

The plasmid superhelical density protocol was adapted from the one described by Folarin *et al*., 2019 (Folarin et al., 2019). Super-helical density was determined by analysing 1.5 μg of pDNA on chloroquine agarose gel electrophoresis after treatment with Topoisomerase I from *E. coli* (BIOKÉ), The Netherlands) during 30 min at 37 °C. Agarose (0.7% w/v) was prepared in 1X TBE (tris-borate-EDTA) buffer. Before running, 2.0 mg/L of chloroquine diphosphate was used to pre-stain the gels. Samples were loaded, and electrophoresis was carried out in the first dimension at 2 V/cm for 24 h for the plasmids pAAV-RC2 and pAAV-ImV and for 30h for the plasmid pAAV-MccV, with 1X TBE containing the same concentration of chloroquine diphosphate as the running buffer. The electrophoresis was stopped, and the gel was allowed to soak in 1X TBE buffer with 6 mg/L chloroquine diphosphate for 3h. Electrophoresis was carried out in the second dimension, 90° to the first at 2 V/cm for 24 h with 1X TBE containing the same concentration of chloroquine diphosphate in the soaking buffer as the running buffer. The gels were then rinsed three times in water for an hour each and later stained with SYBR gold nucleic acid stain (ThermoFisher Scientific)(DNA Stains | Thermo Fisher Scientific - ES, n.d.) for 1 h before visualization under a UV light. The superhelical density was calculated as follows:

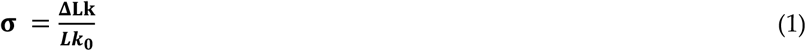

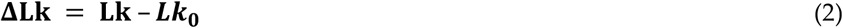

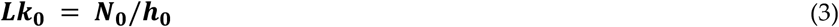

where, *△Lk* is the linking difference determined from the gel image as the number of bands, *Lk*^*0*^ is the number of turns in the relaxed form of plasmid DNA, *Lk* is the number of superhelical turns in pDNA, *N*^*0*^ is plasmid size in base pairs (bp) and *h*^*0*^ are the relaxed base pairs per turns (assumed to be 10.5 bp) (Folarin et al., 2019). The ImageJ software (version 1.53c) (National Institute of Health, n.d.) was used as a densitometry tool to quantify the intensity of the bands on the acquired image of the gel.

### 5.6 Agar diffusion test (ADT)

Antimicrobial activity of microcin V (MccV) was assayed by spotting 5 μL of supernatants onto LB agar (1.5%) plates previously seeded with 10^5^ CFU of a fresh overnight culture of the indicator strain *E. coli* STABLE. The cell-free culture supernatants of the relevant strain(s) were obtained by centrifugation at 12,000 *Xg* for 10 min at 4 °C and subsequent filter sterilization The agar plates were incubated overnight (16 h) at 37 °C and growth inhibition of the indicator microorganism was visualized.

## 6. Acknowledgements

We thank Dr. C.P. Barnes (University College London) for kindly providing the wild type producing strain of microcin V, and Dr. Alex Quintero (Syngulon SA) for his critical review of the manuscript. We would also like to acknowledge the Walloon Region support that enabled this research.

