## Supplementary material for "Bacteriocin peer selection for the production of antibiotic selection free biotherapeutic pDNA"

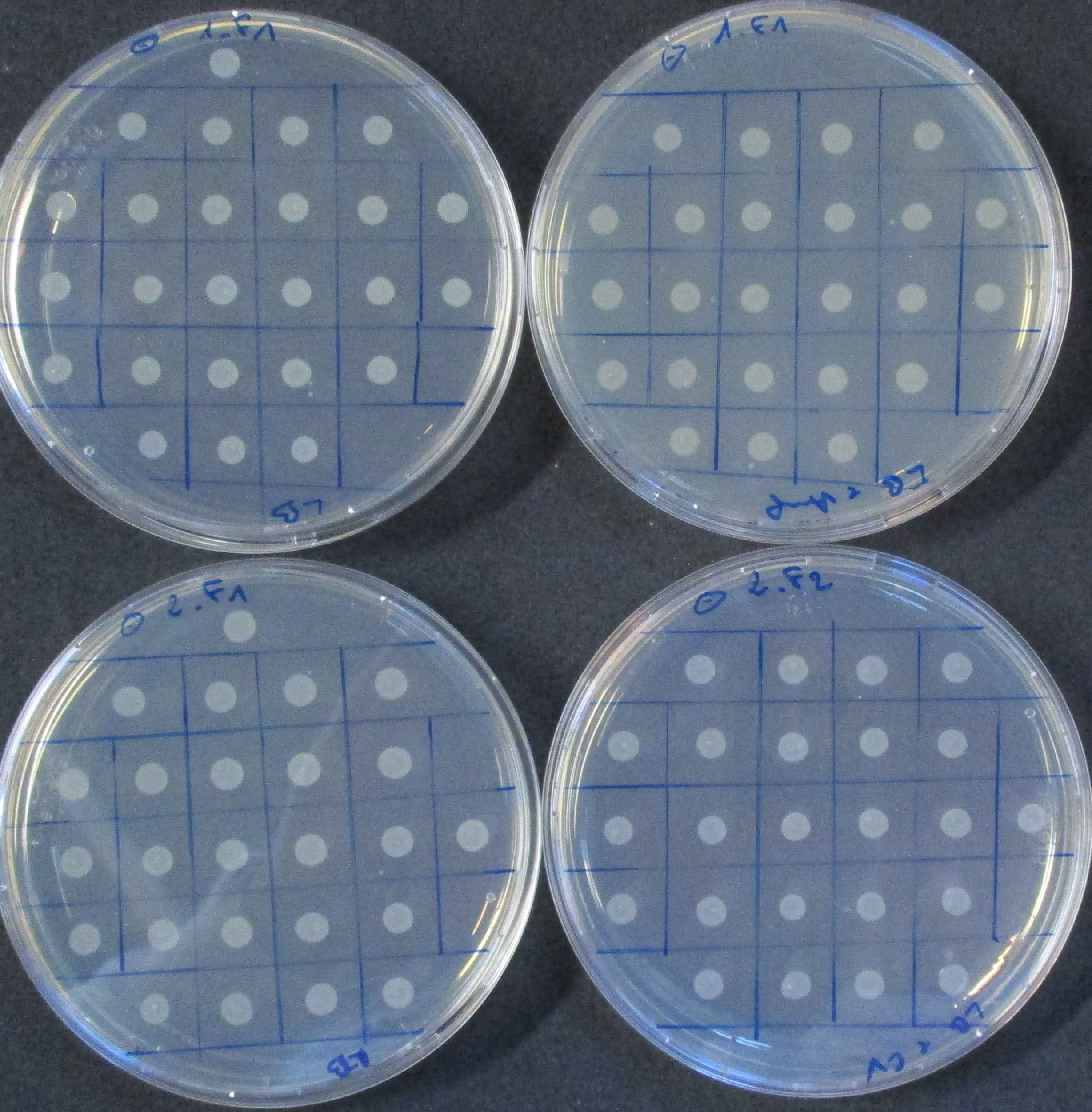

**LB**

**LB + Amp**

**pAAV-RC2**

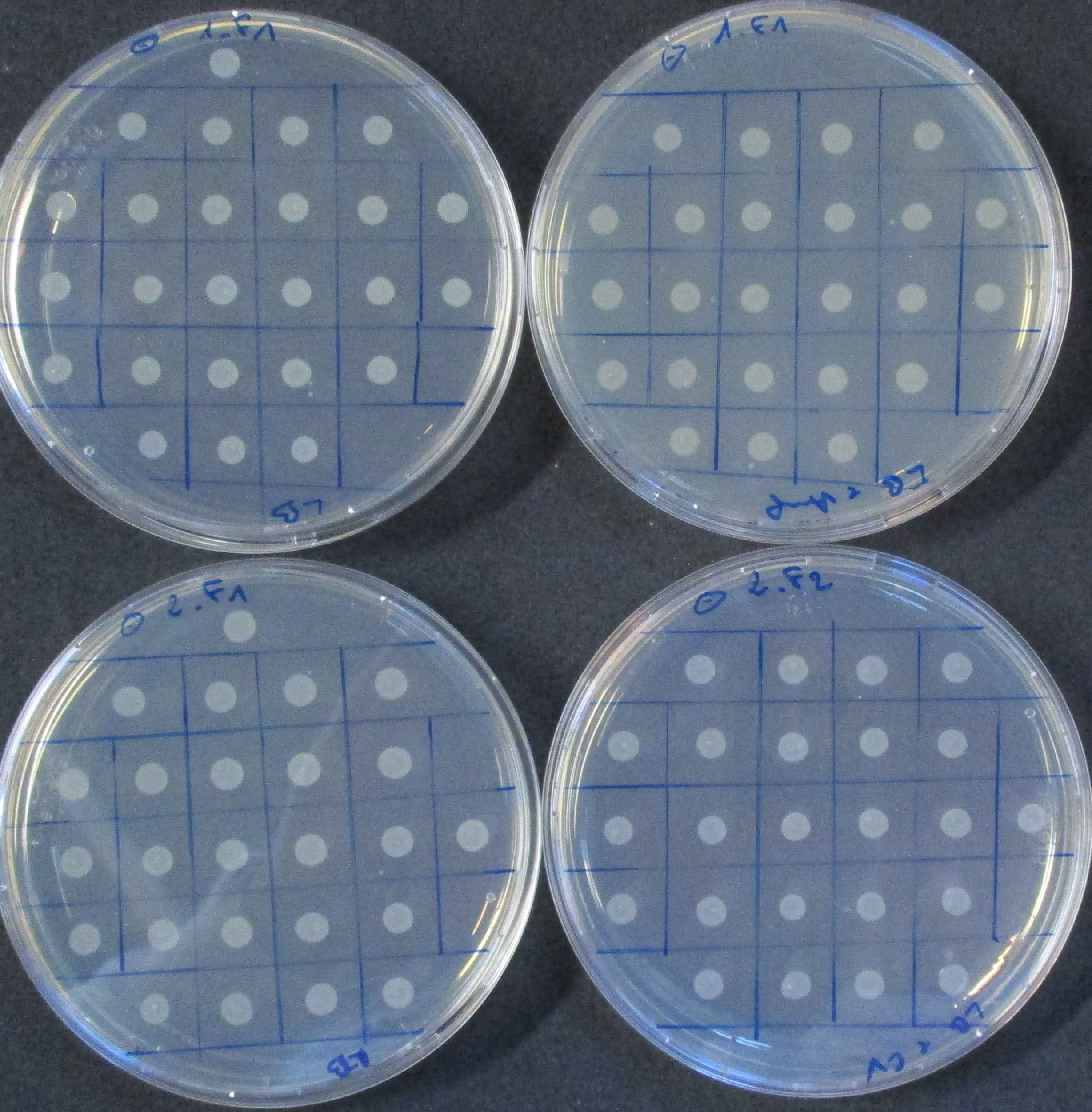

**LB + MccV**

**LB**

**pAAV-MccV**

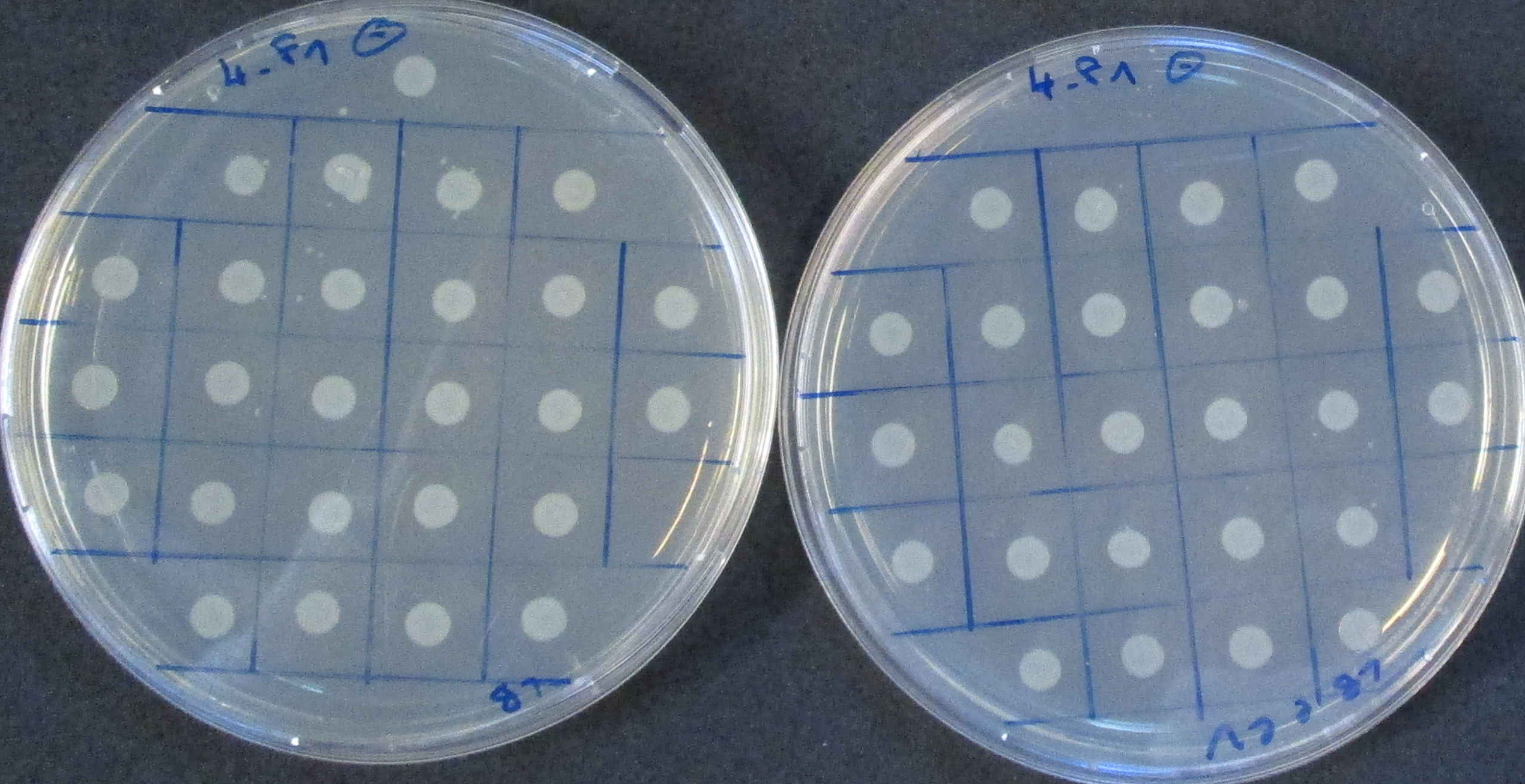

**LB + MccV**

**LB**

**pAAV-ImV**

**-**

**-**

**-**

**Percentage of Stability (%)**

**100 %**

**100 %**

**100 %**

### The stability of each plasmid within its host cell line was verified by growing a *E. coli* STABLE strain with each one of the plasmids used throughout this work (pAAV-RC2, pAAV-MccV and pAAV-ImV) in non-selective (LB) and selective (LB+Kan or LB+MccV) agar plates. The percentage of plasmid retaining cells was determined from the ratio of colonies on selective plates over those on non-selective plates. Negative control (-).
